# A Monoclonal Antibody Panel to Track Antigenic Evolution of SARS-CoV-2 JN.1 Subvariants

**DOI:** 10.64898/2026.05.29.728770

**Authors:** Hsiang Hong, Madeline Wu, Sue Chong, Jian Yu, Ian A. Mellis, Chien-Yu Huang, Qian Wang, Yaoxing Huang, Yicheng Guo, David D. Ho

**Author notes:** Correspondence (Y.G.) & (D.D.H.). These authors contributed equally (Hsiang Hong, Madeline Wu).

## Abstract

SARS-CoV-2 continues to evolve, generating successive variants with enhanced resistance to antibody-mediated neutralization. However, many existing monoclonal antibody (mAb) repertoires were derived from ancestral or early Omicron strains and have lost activity against contemporary variants, limiting their ability to resolve the antigenic effects of mutations arising in newly emerging lineages. Here, we generated a mAb panel by immunizing naïve, unimprinted VelocImmune humanized mice with spike from the KP.3.1.1 variant, and isolated and characterized 11 neutralizing antibodies targeting diverse epitopes of recently evolved antigenic variants. We then integrated these antibodies with previously characterized mAbs to establish an expanded toolbox for profiling antigenic evolution among emerging variants. Application of this panel to currently emerging SARS-CoV-2 variants provided mechanistic insight into recent antigenic evolution, highlighting the potential role of altered receptor-binding domain conformation. Our data also revealed clear differences in antibody-evasion profiles between JN.1-derived lineages and BA.3.2 lineages. Together, our findings establish an updated mAb panel for monitoring antigenic evolution of JN.1 subvariants.

## Introduction

SARS-CoV-2 continues to evolve under widespread population immunity, generating successive variants with enhanced transmissibility and increasing resistance to antibody-mediated neutralization^1,2^. Although the emergence of the JN.1 lineage marked a major antigenic shift^3,4^, continued diversification of its descendants has produced increasingly divergent subvariants harboring additional spike mutations that reshape viral antigenicity^5–8^. Multiple antibody-evasive variants, including KP.3.1.1, XEC, LP.8.1.1, NB.1.8.1, and XFG, have rapidly emerged and competed for regional or global dominance, reflecting continued adaptation under immune selection pressure (**Fig. 1A-B**). This continual antigenic change highlights an ongoing challenge: the need to monitor SARS-CoV-2 immune escape in real time as newly emerging variants accumulate increasingly complex mutational combinations^9^.

**Fig. 1.**
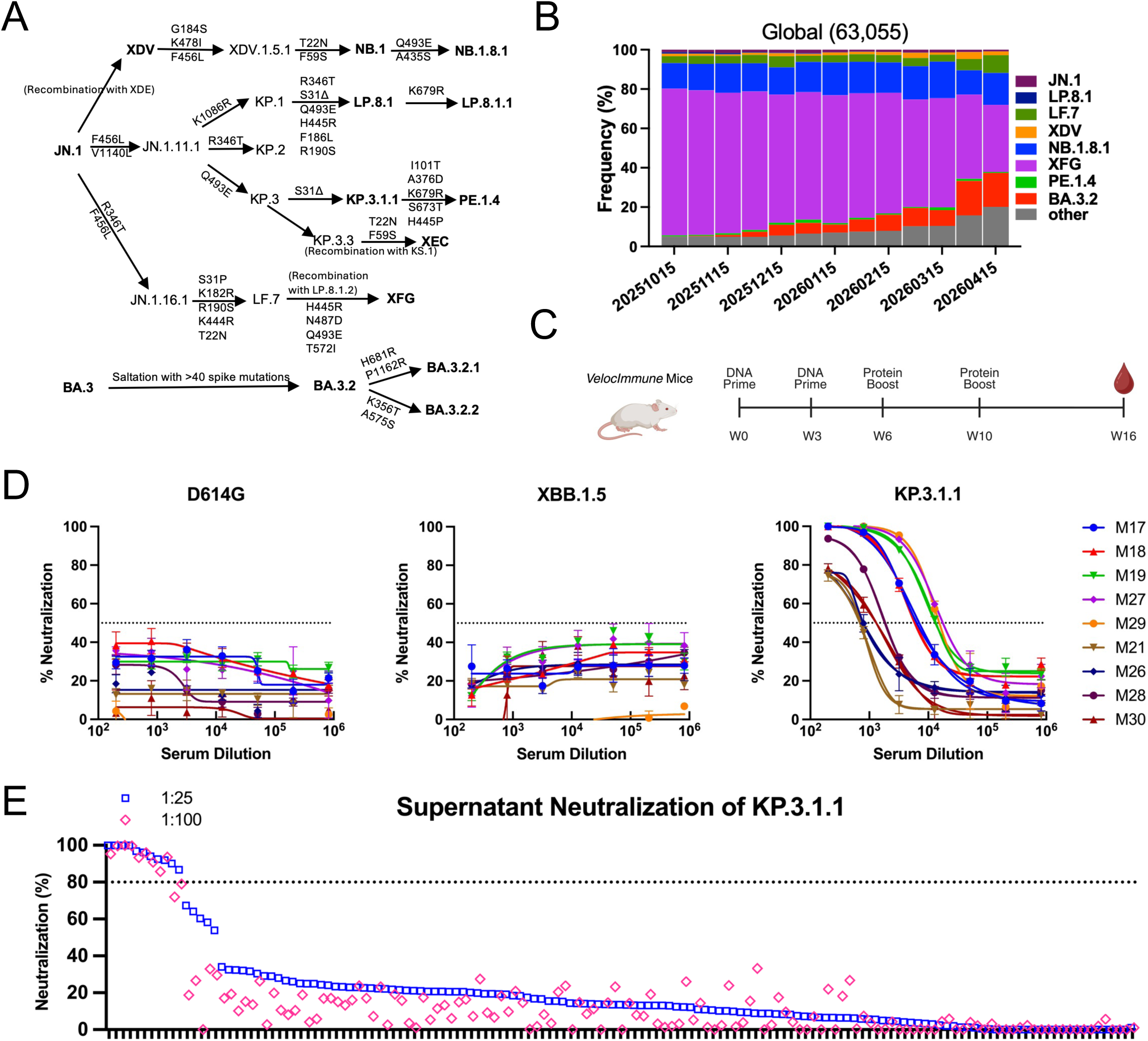
Isolation of neutralizing antibodies against recent SARS-CoV-2 variants from VelocImmune mice. A. Mutation pathway of recent SARS-CoV-2 variants; KP.3.1.1 was chosen as the immunization strain. B. Frequencies of recent variants obtained from October 15, 2025 to April 15, 2026. C. Immunization schedule to isolate neutralizing antibodies. D. Serum neutralization titers against D614G, XBB.1.5, and KP.3.1.1 at week 8. Data are shown as mean ± standard error of mean (s.e.m.) for three technical replicates. E. Supernatant nutralization screening of selected monocloanl antibodies against KP.3.1.1.

Monoclonal antibodies (mAbs) have been instrumental for tracking antigenic evolution, informing therapeutic development, and mapping vulnerable epitopes since the start of the pandemic^10–22^. However, many currently available mAbs were generated against ancestral strains or early Omicron variants and may incompletely represent the antigenic landscape of circulating SARS-CoV-2 variants^23–26^. Progressive accumulation of mutations within the receptor-binding domain (RBD) and N-terminal domain (NTD) has diminished recognition by numerous therapeutic and research antibodies as key neutralizing epitopes continue to diversify^16,17,27^. Beyond amino acid substitutions, recent variants have also exploited structural mechanisms of immune evasion, including altered receptor-binding domain conformational dynamics that reduce antibody accessibility^28,29^. Taken together, these changes raise the possibility that existing monoclonal antibody repertoires are increasingly mismatched to recently circulating strains.

Interpretation of emerging antigenic evolution is further complicated by immune imprinting. Repeated vaccination^30,31^ and infection^32,33^ shape humoral immunity through preferential recall of memory B cell responses against previously encountered antigenic sites, potentially constraining the diversity of antibodies elicited against newly evolved spike proteins^34,35^. Consequently, many human-derived mAbs may disproportionately reflect historical antigenic exposure and may be biased toward epitopes that are differentially mutable or conformationally occluded in circulating variants. At the same time, a small but growing fraction of the population, particularly young children whose earliest SARS-CoV-2 exposures involved recent variants, may have limited or no ancestral-strain imprinting and may therefore develop antibody specificities distinct from those of repeatedly exposed individuals^36,37^. Studying antibody responses generated without prior immune imprinting may provide complementary insight into antigenic features relevant to these minimally imprinted populations, although such responses should not be considered representative of population-level human immunity. Existing antibody panels therefore face an increasing limitation for antigenic surveillance: as their activity is progressively eroded by viral evolution, they provide diminishing resolution for determining the consequences of mutations arising on evolving spike backgrounds. Periodically generating antibodies against recent variants may replenish this toolbox to recognize currently circulating antigenic surfaces, enabling more granular characterization of subsequent viral evolution.

Here, we generated a human monoclonal antibody panel by immunizing naïve VelocImmune mice with the KP.3.1.1 spike protein and isolating neutralizing antibodies against recently circulating SARS-CoV-2 variants. Through functional epitope and neutralization profiling, we established an expanded framework for monitoring antigenic evolution in JN.1 sublineages and the recent saltation variant, BA.3.2. Application of this panel captured the antigenic footprint of JN.1-derived sublineages and the antibody-evasion trajectory of BA.3.2 variants. Notably, this antibody panel revealed functionally convergent mutations associated with enhanced antibody escape in recent subvariants, suggesting that altered spike conformational dynamics may contribute to immune evasion across multiple evolving SARS-CoV-2 variants.

## Results

### Isolation of neutralizing antibodies against recent SARS-CoV-2 variants

To generate human monoclonal antibodies against recent SARS-CoV-2 variants without prior immune imprinting, we immunized VelocImmune mice, which carry human immunoglobulin variable-region genes, with spike from KP.3.1.1, the dominant circulating strain at the time of immunization. Mice received two intramuscular DNA/electroporation-based vaccine doses with plasmids encoding the full-length KP.3.1.1 spike as the primary series, followed by two intramuscular protein-based booster doses using recombinant KP.3.1.1 spike trimer, over a total of 16 weeks (**Fig. 1C**)^38^. We reasoned that immunization with a recent spike antigen in immunologically naïve mice may elicit antibodies targeting antigenic features preferentially exposed in recently circulating variants rather than epitopes shaped by historical SARS-CoV-2 exposure.

Serum neutralizing activity against representative SARS-CoV-2 pseudoviruses were assessed two weeks after each immunization. By week 8, sera from all immunized mice demonstrated robust neutralization against the homologous KP.3.1.1 strain, while exhibiting minimal activities against the ancestral D614G and earlier XBB.1.5 variants (**Fig. 1D**). This restricted neutralization profile suggested successful elicitation of antibody responses preferentially directed toward recently evolved spike antigenic sites.

To isolate mAbs mediating these responses, five mice (ID#: 17, 18, 19, 27, 29) with the highest serum neutralization titers were selected for antigen-specific B-cell sorting^38^. Splenocytes and peripheral blood mononuclear cells (PBMCs) were stained with a fluorescently labeled KP.3.1.1 spike trimer, and antigen-specific B cells were flow sorted using a panel of established cell surface marker-directed antibodies. Next, we conducted 10X Genomics single-cell sequencing of sorted B cells to obtain paired heavy- and light-chain variable sequences. From 12,253 B-cell clones with paired heavy- and light-chain BCR sequences, we selected 144 candidate mAbs based on cell type, isotype, somatic hypermutation level, CDR3 length, germline gene utilization, and clonal expansion (**Fig S1A-B**). Following cloning and expression, these candidates were screened them for spike binding and neutralizing activity. Of the 144 mAbs, 77 of them could bind to KP.3.1.1 spike (**Fig. S1C**), 11 achieved ≥80% inhibition of KP.3.1.1 pseudovirus at a 1:25 culture supernatant dilution and were selected as the primary antibody candidates (hereafter referred to as the C-series mAbs with identifiers starting with “C”) for downstream characterization (**Fig. 1E**).

### Functional epitope assignment for C-series mAbs

To define the antigenic landscape recognized by the C-series antibodies, we first assessed binding against the KP.3.1.1 recombinant spike trimer, RBD, and NTD antigens by ELISA (**Fig. 2A**). All 11 mAbs bound to full-length spike protein, with eight antibodies (C46, C52, C67, C104, C116, C118, C129, and C136) demonstrating selective binding to the RBD, while three antibodies (C27, C94, and C131) bound the NTD. These findings indicate that the C-series panel spans multiple major surfaces on spike and is not restricted to a single antigenic region.

**Fig. 2.**
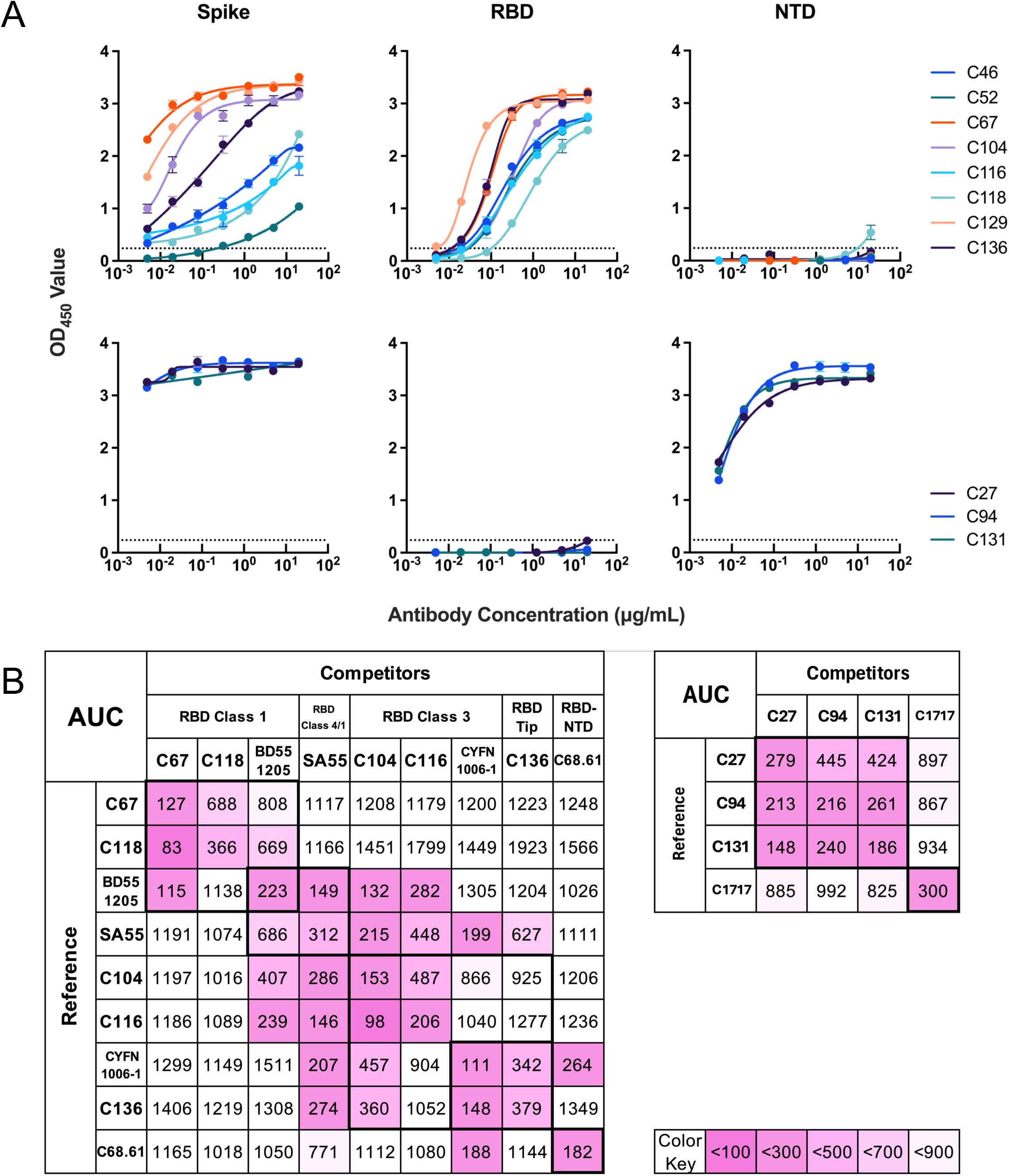
Functional epitope profiling of the C-series mAbs. A. ELISA binding of KP.3.1.1 spike, RBD, and NTD by purified mAbs. Data are shown as mean ± standard error of mean (s.e.m.) for three technical replicates. B. Competition ELISA matrix showing the ability of unlabeled spike-binding mAbs to block binding of a biotinylated mAb to the spike trimer. Values denote the area under the competition curve (AUC). Color denotes competition level, with darker colors corresponding to more competition.

To further define epitope relationships, we performed competition ELISAs among most C-series mAbs and against several previously characterized reference monoclonal antibodies^39–42^ that retained neutralizing activities against recent variants with known SARS-CoV-2 antigenic sites (**Fig. 2B**). Among RBD-targeting antibodies, C67 and C118 competed with one another and with the class 1 antibody BD55-1205^40^, consistent with recognition of class 1-like epitopes. C104 and C116 formed a distinct competition cluster with the class 3 antibody CYFN1006-1^39^, suggesting recognition of class 3-like epitopes. The NTD-targeting antibodies C27, C94, and C131 also formed a separate competition cluster distinct from the previously characterized NTD antibody C1717^42^, suggesting recognition of related but non-identical antigenic surfaces. Due to weak RBD binding, C46, C52, and C129 were not resolved by the competition matrix and were assigned using epitope-directed mutagenesis.

To further understand and validate the functional epitope profiles of our mAbs, we introduced mutations disrupting spike epitopes (**Fig. S2A**) and assessed antibody binding by ELISAs (**Fig. S2B**). Mutations within the RBD class 1 epitope (N417A, S455A, Q493E, and H505A) dramatically reduced spike binding of C67 and C118, supporting their class 1-like epitope assignment. Likewise, mutations within the RBD class 3 epitope (N437R, K440A, L441R, K444A, H445R) substantially reduced binding of C104 and C116. For C129, mutations in the RBD class 4 epitope (G504D, H505A) knocked out its binding to the spike trimer, supporting recognition of a class 4-like epitope. C46, C52, and C136 exhibited reduced binding to mutations introduced in the RBD tip region (K478I, N477R, P486A), indicating targeting of this antigenic surface (**Fig. S2B**). Among NTD-targeting antibodies, NTD supersite mutations abolished binding of C94 and C131, whereas C27 retained detectable binding, suggesting recognition of a neighboring or partially overlapping NTD epitope distinct from the canonical supersite.

To assess whether the binding reductions observed for the class 3 and RBD-tip mutants reflected epitope-specific disruption rather than broader effects on antigen integrity, we cross-tested these mutants against antibodies targeting other antigenic regions (**Fig. S3**). The class 3 mutant RBD selectively reduced binding of class 3-directed antibodies C104 and C116, whereas antibodies assigned to other RBD epitope classes largely retained binding (**Fig. S3A**). Similarly, RBD-tip-directed antibodies C46, C52 and C136 showed reduced binding to the RBD-tip-mutated spike, while antibodies targeting other spike epitopes retained binding (**Fig. S3B**). These cross-testing results further support the functional epitope assignments of the C-series antibodies.

Collectively, these findings establish the C-series antibodies as a diverse panel spanning multiple major antigenic regions of the spike of recent SARS-CoV-2 variants.

### C-series mAbs preferentially neutralize recent SARS-CoV-2 variants

To evaluate neutralization breadth and potency of the C-series mAbs, we tested their neutralizing activities against representative SARS-CoV-2 pseudoviruses spanning ancestral (D614G), Omicron (BA.5), XBB.1.5, and JN.1-descendant variants (JN.1 and KP.3.1.1; **Fig. 3A**). Consistent with the KP.3.1.1-focused serum responses in **Fig. 1D** observed following immunization, all antibodies demonstrated potent neutralizing activity against the homologous KP.3.1.1 strain and its parental JN.1 lineage, with IC50 values largely within the nM to pM range (**Fig. 3B**).

**Fig. 3.**
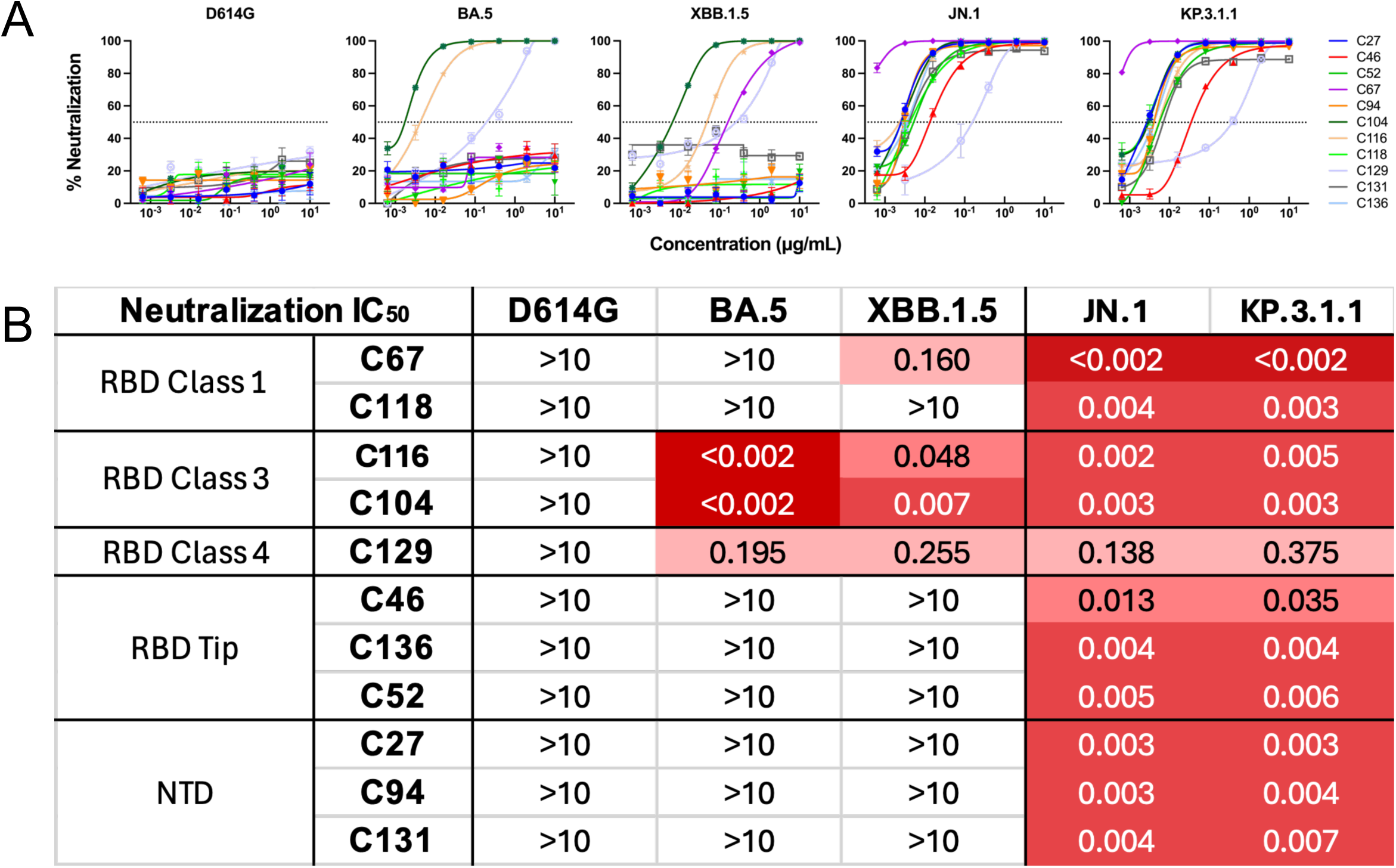
Neutralization profiles of the C-series mAbs. A. Neutralization of select SARS-CoV-2 pseudoviruses by purified mAbs. Data are shown as mean ± standard error of mean (s.e.m.) for three technical replicates. B. Neutralization IC50 values for the 11 mAbs, grouped by epitope classes.

In contrast, neutralizing activity against earlier SARS-CoV-2 variants was substantially restricted. None of the antibodies demonstrated appreciable activity against ancestral D614G, highlighting the antigenic divergence between historical and recent spike proteins. Despite this restricted breadth, selective cross-neutralization was observed against earlier Omicron variants. Notably, three antibodies retained activity against BA.5, with the class 3 antibodies C104 and C116 maintaining high potency despite the substantial antigenic distance between BA.5 and the KP.3.1.1 immunogen^6^. Similarly, four antibodies demonstrated measurable neutralization of XBB.1.5, indicating partial conservation of select antigenic surfaces across successive Omicron waves. Together, these findings demonstrate that the C-series antibodies preferentially recognize antigenic features enriched in recently circulating SARS-CoV-2 variants rather than broadly conserved ancestral determinants, supporting their utility for monitoring ongoing antigenic evolution in a lineage divergent from older strains.

### An expanded antibody panel reveals enhanced escape of the recent SARS-CoV-2 variants

As SARS-CoV-2 continued to diversify during this study (**Fig 1A**), XFG became the dominant variant in mid-2025^8^, whereas BA.3.2 has recently increased gradually at the global level (**Fig. 1B**), with substantially higher frequencies observed in Europe and Oceania (**Fig. S4**). In parallel, PE.1.4, a KP.3.1.1 sublineage, rose to a noticeable frequency in Oceania since October of 2025, where variant circulation patterns differed from those in Europe and North America. To characterize the antigenic consequences of these evolutionary trajectories, we expanded our antibody panel to assess the antigenic escape of newly emerging variants. We first assembled a 23-mAb panel comprised of the newly generated C-series antibodies, together with previously characterized and available mAbs that retained activity against recent variants targeting distinct epitopes ^24–26,39–44^ (**Fig. 4A**).

**Fig. 4.**
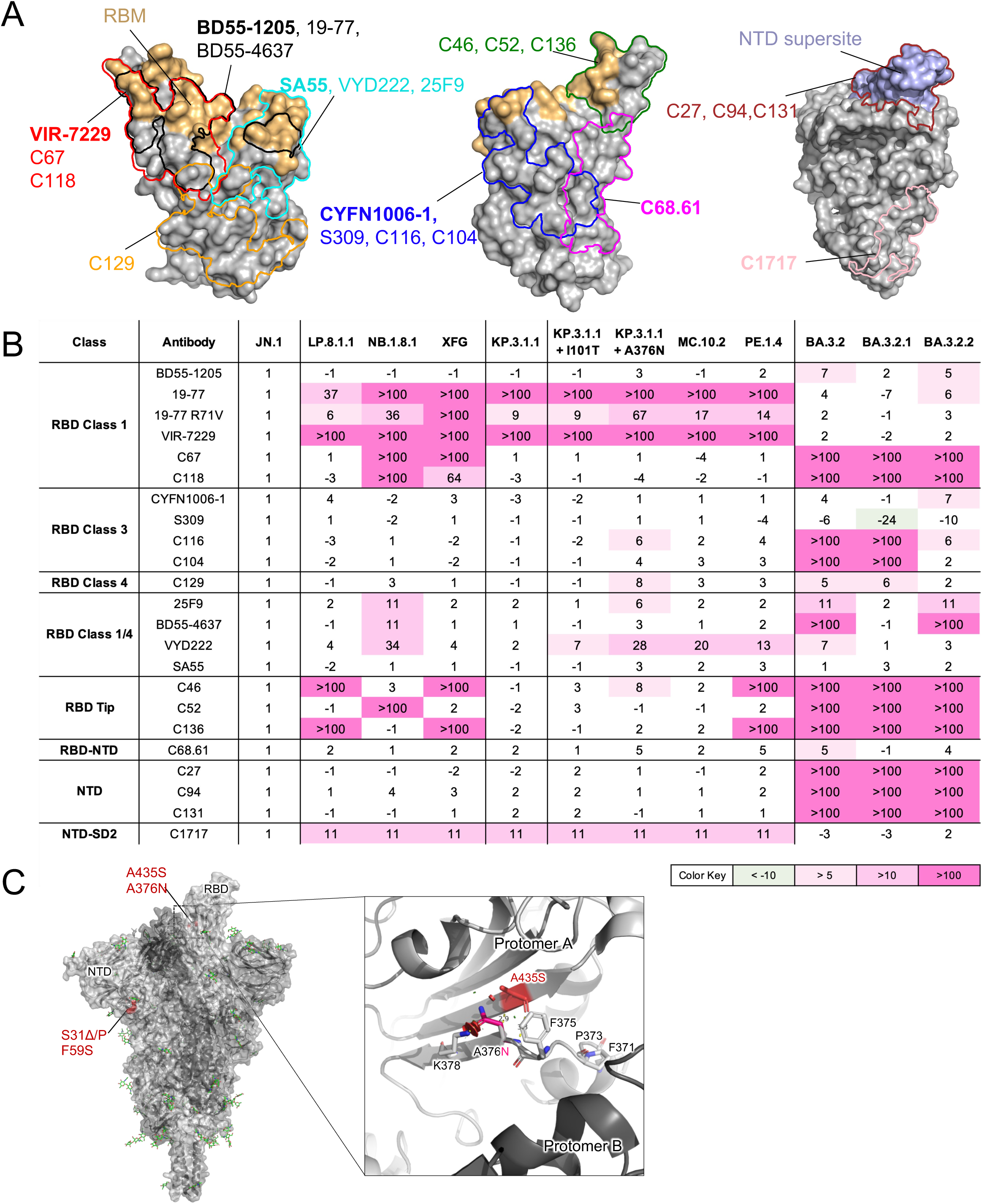
Antigenic tracking of recent SARS-CoV-2 variants. A. Antibody panel used to track antigenic variation among SARS-CoV-2 variants. Epitope footprints are shown for the representative mAbs indicated in bold. The hACE2-binding site is shown in light orange, and the NTD supersite is shown in light blue. B. Fold change in neutralization IC_50_ values relative to JN.1. C. Structural model showing the location of A376N, found in PE.1.4, relative to selected mutations present in KP.3.1.1, NB.1.8.1, and XFG.

We next tested this enhanced mAb panel for neutralizing activity against a set of 13 circulating and historical SARS-CoV-2 variants, including JN.1 and its descendant sublineages LP.8.1.1, NB.1.8.1, XFG, and the homologous KP.3.1.1 strain (**Fig. 4B & S5**). Compared with the parental JN.1 lineage, most circulating variants retained broad susceptibility to NTD-directed mAbs, whereas RBD-directed mAbs exhibited variant-specific escape patterns. LP.8.1.1 and XFG showed reduced sensitivity to primarily class 1 antibodies, with more modest escape from select class 1/4, and class 3 antibodies, while NB.1.8.1 primarily escaped class 1 and class 1/4 antibodies. These results are consistent with previous observations^8^ but more extensive with higher-resolution epitope-level mapping across our expanded antibody panel. In addition, PE.1.4, which has intermittently increased in frequency in Oceania, demonstrated broader reductions in susceptibility to neutralization across multiple antibodies. Antibody escape was particularly pronounced among class 3-directed antibodies, with 7-fold increase in IC50 relative to its parental KP.3.1.1 strain by C116 and complete loss of detectable neutralization by C136, suggesting epitope-specific patterns of antibody evasion among currently circulating JN.1 subvariants.

PE.1.4 contains five additional spike substitutions relative to its parental KP.3.1.1 lineage (**Fig. 1A**), including I101T within the NTD and A376N within the RBD. To assess the contribution of these substitutions to antibody escape, we generated pseudoviruses encoding KP.3.1.1 carrying I101T, A376N, or both substitutions (also known as the subvariant MC.10.2). Relative to KP.3.1.1, introduction of A376N alone resulted in broad reductions in neutralization sensitivity across multiple antibodies, particularly those sensitive to spike conformational changes, including RBD class 1, class 3, class 1/4 antibodies, as well as the RBD-NTD interface mAb C68.61^41^ (**Fig. 4B & S5**). In contrast, introduction of I101T had no detectable effect on neutralization across the antibody panel. The combined mutant MC.10.2 partially recapitulated the PE.1.4 neutralization phenotype, indicating that A376N is a major contributor to the enhanced antibody escape observed in PE.1.4.

We additionally evaluated the expanded antibody panel against the highly divergent saltation variant BA.3.2, which contains more than 50 additional spike substitutions relative to its parental BA.3 lineage and is both genetically and antigenically distant from contemporary JN.1-descendant variants^5,7,45^. None of the C-series antibodies demonstrated neutralizing activity against BA.3.2, consistent with their known preferential recognition of JN.1-derived antigenic features compared to earlier more dominant strains. In contrast, several previously characterized broadly neutralizing antibodies maintained measurable neutralizing activity against BA.3.2, especially for the class 1 antibodies, perhaps partially explaining why despite the substantial antigenic divergence from currently circulating strains, the strain has yet to become dominant.

### A376N may promote antibody escape through altered spike conformational dynamics

The sequential replacement of PE.1.4 by XFG and subsequently by BA.3.2 in Oceania provides insight into the mechanisms shaping current SARS-CoV-2 evolution. Given the prominent contribution of A376N to the antibody-evasion phenotype of PE.1.4, we investigated how this mutation affects neutralization sensitivity. A376N lies within the SA55 epitope, providing a direct structural explanation for the approximately threefold reduction in SA55 neutralization caused by this mutation (**Fig. S6A**). In contrast, A376N is located far from the binding sites of other broadly neutralizing class 1 mAbs, including 19-77 and VIR-7229, suggesting that its effects on these antibodies are mediated indirectly through long-range conformational changes rather than direct disruption of antibody–RBD contacts (**Fig. S6A**). A similar allosteric effect was observed for the conformation-sensitive mAb C68.61, which targets the outer face of the RBD, whereas residue 376 is located on the inner face. Consistent with this interpretation, A376N also reduced sensitivity to hACE2-mediated inhibition by approximately 4-fold despite being spatially distant from the hACE2-binding interface, further supporting that this mutation favors the RBD-down conformation and thereby limits hACE2 access to the receptor-binding site (**Fig. S6B**). Further structural analysis revealed that A376N lies adjacent to both A435S and the 371–375 loop, regions positioned near the interprotomer interface between neighboring RBDs (**Fig. 4C**). The 371–375 loop has been linked to RBD conformational regulation and has remained fixed since the emergence of Omicron sublineages^16,17^. In parallel, recent variants such as XEC and XFG harbor S31 and F59 mutations that may also modulate RBD conformational dynamics^8,27^. Moreover, A435S, present in NB.1.8.1 and BA.3.2, has been associated with altered RBD conformation^8^, the BA.3.2.1 has been shown in predominantly close conformation^46^. Together, these findings suggest that A376N not only could mediate antibody evasion by alter the epitopes of antibodies target the RBD inner face but also may provide an additional route for stabilizing interactions that favor the RBD-down state, thereby reducing exposure of neutralization-sensitive epitopes.

## Discussion

As SARS-CoV-2 continues to evolve under immune selective pressure, tracking antigenic evolution in real time is increasingly challenging^1,9^. In this study, we generated a panel of human monoclonal antibodies independent of ancestral immune imprinting through immunization of naïve VelocImmune mice with the KP.3.1.1 spike and characterization of their neutralization breadth against recent SARS-CoV-2 variants. The resulting C-series mAbs preferentially and robustly neutralized the dominant JN.1-descendants while limited activity against earlier ancestral and pre-JN.1 strains, consistent with recognition of antigenic features enriched in recently circulating viruses. Together with previously characterized antibodies, the expanded panel provided a useful toolbox for profiling antigenic changes among newly emerging variants.

Functional epitope profiling demonstrated that the C-series mAbs span multiple neutralizing surfaces across the RBD and NTD. Although many currently available monoclonal antibodies were generated against ancestral or early Omicron strains, the continued diversification of SARS-CoV-2 has progressively altered vulnerable antigenic surfaces. The restricted breadth of the small, potency-selected C-series panel is consistent with the possibility that some antigenic features of recent spikes differ from those recognized by antibodies elicited by earlier SARS-CoV-2 strains. However, larger and more representative antibody collections will be required to determine how broadly this observation applies. It is also worthwhile to note that many of the previously isolated mAbs from ancestral or earlier variants were class 2 antibodies^10,47^, but none was identified in this study, further hinting at the substantial differences in antigenic features of recent variants. In this context, periodic updating of antibody panels may facilitate improved resolution of ongoing antigenic evolution. Future studies of the structural basis of spike neutralization by C-series and similarly raised mAbs will be necessary to specifically define their epitopes and how they differ from competing antibodies.

The expanded antibody panel provided additional resolution for comparing antigenic changes among contemporary SARS-CoV-2 variants, particularly JN.1-derived lineages. Together with epidemiological surveillance, our data characterized the antigenic changes associated with the emergence of PE.1.4 and its subsequent replacement by XFG and BA.3.2. A376N was a major contributor to the broad antibody-evasion phenotype of PE.1.4. Its effects on conformation-sensitive class 1 and class 1/4 mAbs, together with reduced hACE2-binding, are consistent with the possibility that A376N alters RBD accessibility or conformational dynamics. However, because the region surrounding A376N is known to be immunodominant^48–50^ and mutations at adjacent residues have been reported to alter multiple spike and viral phenotypes^51–54^,direct epitope disruption and other functional effects cannot be excluded. Therefore, direct structural or biophysical evidence will be required to determine whether it favors a more closed, RBD-down state. XFG and BA.3.2 contain distinct mutations in the NTD and RBD, respectively, that may similarly affect spike conformation or antibody accessibility, while additional mutations could provide further antigenic or fitness advantages.

The highly divergent BA.3.2 lineage escaped nearly all JN.1-specific mAbs in our panel but remained susceptible to several broadly neutralizing class 1 antibodies. This residual susceptibility may have constrained its population-level expansion, although viral fitness and the heterogeneous immune histories of human populations are also likely to contribute. Consistent with the importance of exposure history, our recent study showed that sera from young children exhibited neutralization profiles distinct from those of ancestrally imprinted adults, including markedly reduced activity against BA.3.2.2^36^. This finding qualitatively parallels the restricted activity of the C-series mAbs against BA.3.2-related variants but does not establish that antibodies generated in naïve mice recapitulate pediatric human immunity. The specificity of our KP.3.1.1 elicit antibodies to the dominant circulating variants enabled resolution of antigenic differences among recently circulating SARS-CoV-2 variants and identified disproportionate escape in PE.1.4. Continued updating of monoclonal antibody repertoires may therefore remain important for monitoring the antigenic consequences of SARS-CoV-2 evolution as new lineages emerge.

### Limitation of this study

The C-series antibodies were generated in naïve mice and may not reflect the repertoires of most contemporary humans with complex vaccination and infection histories. Moreover, the 11 antibodies examined represent a small, potency-selected subset of the recovered candidates, with only three targeting the NTD, limiting conclusions about epitope immunodominance and NTD antigenic drift. Epitope assignments and the proposed effect of A376N on RBD accessibility were based on functional evidence; structural and biophysical studies will be required to define the binding interfaces and underlying mechanisms.

## STAR METHODS

Detailed methods are provided in the online version of this paper and include the following:

### RESOURCE AVAILABILITY

#### Lead contact

Further information and requests for resources and reagents should be directed to and will be fulfilled by the Lead Contact Author David D. Ho.

#### Materials availability

All reagents generated in this study are available from the Lead Contact with a completed Materials Transfer Agreement.

#### Data and code availability

Data reported in this paper will be shared by the lead contact upon request. This paper does not report original code. The C-series antibody sequences were deposited in GenBank, under accession: PZ745502 to PZ745523. Any additional information required to reanalyze the data reported in this paper is available from the lead contact upon request.

### EXPERIMENTAL MODEL AND SUBJECT DETAILS

#### Cell Lines

HEK293T (CRL-3216) and Vero-E6 (CRL-1586) cells were purchased from ATCC and cultured at 37°C with 5% CO2 in Dulbecco’s Modified Eagle Medium (DMEM) + 10% fetal bovine serum (FBS) + 1% penicillin-streptomycin. Expi293 cells (A14527) were purchased from Thermo Fisher Scientific and maintained in Expi293 expression medium per the manufacturer’s instructions. The morphology of each cell line was confirmed visually before use. All cell lines tested mycoplasma negative. Vero-E6 are from African green monkey kidneys. HEK293T and Expi293 cells are of human female origin.

### METHOD DETAILS

#### SARS-CoV-2 spike plasmids

SARS-CoV-2 spike-expressing plasmids were generated using previously reported methods^26^. To generate the expression constructs for soluble spike trimer (S2P) proteins, the ectodomains (1-1208aa, numbering based on WA1) of the spikes were PCR amplified and cloned into the paH vector and then introduced K986P and V987P substitutions, as well as a “GSAS” substitution of the furin cleavage site (682-685aa in WA1) into the spikes. KP.3.1.1 S2P was fused with a Strep-Tag II at the C terminus, followed by a FLAG tag. All constructs were confirmed by whole-plasmid sequencing prior to pseudovirus packaging.

Antibody-expressing constructs were generated as previously described^26^. The variable regions of heavy and light chains for each antibody were synthesized (Twist Bioscience) and then cloned into the gWiz vector. To make spike/antibody plasmid constructs carrying individual mutations, the QuikChange Site-Directed Mutagenesis Kit (Agilent) was utilized following the manufacturer’s instructions.

#### Protein expression and purification

Plasmid was transfected into Expi293 cells using 1 mg/mL PEI-MAX at a ratio of 1:3, and the supernatants were collected after 5 days. Purification of cell supernatant was performed using

rProtein A Sepharose (Cytiva). For the spike trimer proteins or RBD proteins, paH-spike or p3BNC-RBD, respectively, was transfected into Expi293 cells using PEI-MAX at a ratio of 1:3, and the supernatants were collected five days later. The His-tagged and the Strep-tagged proteins were purified using Excel resin (Cytiva) and Strep-Tactin^®^XT 4Flow^®^ (IBA Lifesciences) following the manufacturer’s instructions. Molecular weight and purity were confirmed by SDS-PAGE protein electrophoresis and size-exclusion chromatography prior to use.

#### Mouse vaccination

VelocImmune female mice were acquired from the manufacturer (Regeneron Pharmaceuticals, Inc.) and immunized with 10 µg each of KP.3.1.1-expressing plasmids delivered into both hind limbs by DNA electroporation at weeks 0 and 3. Protein boosts (10 µg of purified soluble spike trimer per dose) were administered intramuscularly at weeks 6 and 10. Peripheral blood was collected two weeks after each immunization to monitor serum ELISA binding and neutralization titers. Splenocytes and peripheral blood mononuclear cells (PBMCs) were isolated at week 16 for B-cell sorting and downstream antibody characterization. All animal procedures were reviewed and approved by the Institutional Animal Care and Use Committee of Columbia University.

#### Antigen-specific memory B cell sorting and single-cell B cell receptor sequencing

Splenocytes and PBMCs were stained with the LIVE/DEAD™ Fixable Yellow Dead Cell Stain Kit (Invitrogen) at ambient temperature for 10 min, washed with RPMI-1640 complete medium supplemented with 2% FBS, and incubated with 10 µg/ml of KP.3.1.1 S2P at ambient temperature for 45 min. Afterwards, cells were washed again and stained at 4°C for 45 min with a cocktail of flow-cytometry antibodies, including CD3 PerCP-Cy5.5 (BioLegend), CD19 PE (BioLegend), B220 APC-H7 (BD Biosciences), anti-FLAG BV421 (BioLegend), and anti-Strep-TagII APC (IBA). Stained cells were washed, resuspended in RPMI-1640 + 2% FBS, and sorted for antigen-specific memory B cells (CD3⁻ CD19⁺ B220⁺ antigen⁺ live single lymphocytes) by flow cytometry.

Sorted cells were loaded onto a Chromium X controller using the 5′ Single Cell Immune Profiling Assay (10x Genomics) at the Columbia University Genome Center. Library preparation and quality-control steps were performed according to the manufacturer’s instructions, and sequencing was carried out on an Illumina NextSeq 500 platform.

#### Single cell gene expression analysis and identification of spike specific antibody transcripts

The assembly of full-length antibody and other gene transcripts was performed utilizing the Cell Ranger V(D)J analysis software (version 3.1.0, 10x Genomics), employing default settings with the GRCh38 genome and V(D)J germline sequence version 2.0.0 as the reference. To determine the cell type of B cells, we utilized Seurat package version 4.1.0. Cells with fewer than 500 unique molecular identifiers (UMIs), more than 4500 UMIs, or expression mitochondria gene fraction of more than 5% were discarded in the following analysis. The UMI counts were normalized to 10,000 counts per cell and converted to log scale (Seurat function NormalizeData). nUMI were regressed out using the ScaleData function. Highly variable genes (n=3000) were computed and used as input for principal components analysis. Significant principal components were used for downstream graph-based, semi-unsupervised clustering into distinct populations (FindClusters function), and Uniform Manifold Approximation and Projection (UMAP) dimensionality reduction was then used to project these populations in two dimensions. For clustering, the resolution parameter, which indirectly controls the number of clusters, was approximated based on the number of cells according to Seurat guidelines; a vector of resolution parameters was passed to the FindClusters function, and the optimal resolution that established discernible clusters with distinct marker gene expression was selected. To identify marker genes, the clusters were compared pairwise for differential gene expression using the Wilcoxon rank sum test for single-cell gene expression (FindAllMarkers function, min.pct = 0.4, min.diff.pct = 0.3, and false discovery rate < 0.05). To assign identities to these subpopulations, we cross-referenced their marker genes with known B cell subtype markers from the literature, Fcer2a for Naïve B cell, Cd80 for memory B cell, Itgax for atypical memory B, Sdc1 and Xbp1 for plasma blast.

#### Antibody transcript annotation

Transcripts specific to the antigen were analyzed and annotated with SONAR version 2.0, following previously established procedures^17^. Assignment of V(D)J gene segments to each transcript was conducted via BLASTn, employing specialized parameters against a germline gene repository sourced from the International ImMunoGeneTics (IMGT) information system database. The identification of the complementarity-determining region 3 (CDR3) utilized BLAST alignments of the V and J segments, focusing on the conserved second cysteine within the V segment and the WGXG (for heavy chains) or FGXG (for light chains) motifs in the J segment, with “X” indicating any amino acid. Isotype determination for heavy chain transcripts was achieved by analyzing Constant domain 1 (CH1) sequences against a human CH1 gene database from IMGT, using BLASTn with standard parameters. The CH1 allele presenting the lowest E-value was selected for precise isotype classification, adhering to a BLAST E-value cutoff of 10e-6.

Transcripts with incomplete V(D)J segments, frameshifts, or extraneous sequences beyond the V(D)J region were discarded. The filtered transcripts were then aligned to their corresponding germline V gene using CLUSTALO, and levels of somatic hypermutation were quantified through the Sievers method. In instances where cells possessed multiple high-quality heavy or light chains, potentially indicative of doublets, combinations of all H and L chains were generated.

#### Antibody expression and purification

For each antibody, variable region genes were codon-optimized for human cell expression and synthesized by Twist Bioscience. The VH and VL genes were cloned separately into gWiz mammalian expression vectors encoding the respective human IgG heavy- and light-chain constant regions. Monoclonal antibodies were expressed in Expi293 cells by co-transfection of the heavy- and light-chain plasmids using PEI-MAX (1 mg/mL) and cultured at 37°C shaking at 125 rpm in 8% CO2. At day 3 post-transfection, culture supernatant was collected for screening antibody binding by ELISA, and at day 4, additional supernatant was collected for pseudovirus neutralization assays. The remaining culture supernatants were harvested on day 5 for purification using rProtein A Sepharose (Cytiva) affinity chromatography per the manufacturer’s instructions.

#### ELISA

High-binding 96-well plates (Corning Costar) were coated with 50ng per well of the respective purified SARS-CoV-2 trimer or RBD in PBS and incubated at 4°C overnight. Plates were washed with PBST (0.05% Tween-20 in PBS), plates were blocked with 200µL of blocking buffer (1% BSA in PBS) at 37°C for 2 h. After an additional wash step, antibody-containing supernatants or purified monoclonal antibodies were serially diluted in blocking buffer, added to plates, and incubated for 1 h at 37°C. After an additional wash, 100 µL of horseradish peroxidase (HRP)-conjugated goat anti-human IgG (H+L) secondary antibody (Jackson ImmunoResearch; 1:5,000 dilution) was added and incubated for 1 h at 37°C. Following a final wash, Ultra TMB-ELISA substrate solution (ThermoFisher) was added, and the reaction was stopped with 1M sulfuric acid. Absorbance was measured at 450 nm using a microplate reader.

#### Competition ELISA

Competition ELISAs were performed as previously described^38^. Purified monoclonal antibodies were biotinylated using the One-Step Antibody Biotinylation Kit (Miltenyi Biotec) according to the manufacturer’s instructions and subsequently purified with a 50 kDa Amicon^®^ Ultra centrifugal filter (Sigma). Serially diluted competitor antibodies (50 µL per well) were added to purified, respective SARS-CoV-2 trimer or RBD coated ELISA plates, followed by 50 µL of biotinylated antibodies at a concentration that yielded an OD450 value of approximately 1.0 in the absence of competitor antibodies. Plates were incubated at 37°C for 1 h and washed with PBST (0.05% Tween-20 in PBS). Subsequently, plates were incubated for another 1 h at 37°C with 100 µL of Pierce™ High Sensitivity NeutrAvidin™-HRP (ThermoFisher) diluted 1:10,000. After an additional wash, Ultra TMB-ELISA substrate solution (ThermoFisher) was added, and the reaction was stopped with 1M sulfuric acid. Absorbance was measured at 450 nm using a microplate reader. For all competition ELISA experiments, the relative binding of biotinylated antibodies to the SARS-CoV-2 protein in the presence of competitors was normalized to the signal obtained in competitor-free wells. Relative binding curves and area under the curve (AUC) values were determined by fitting non-linear five-parameter logistic (5PL) dose-response models in GraphPad Prism v10.3.1.

#### SARS-CoV-2 pseudovirus packaging

Pseudotyped SARS-CoV-2 was produced following a previously established protocol^26^. HEK293T cells were first transfected with spike-encoding plasmids using 1 mg/mL of PEI-MAX (Polysciences, Inc.) and cultured for 24 hours. The transfected HEK293T cells were then infected with VSV-G pseudotyped ΔG-luciferase virus (Kerafast, EH1020-PM) at a multiplicity of infection (MOI) of approximately 3 to 5. Two hours later, the cells were washed three times and cultured in fresh cell medium overnight. The virus was then harvested first by centrifugation at 2000 rpm for 10 minutes followed by collection of the supernatant. 20% of I1 hybridoma (ATCC, CRL-2700) supernatant was then added to each virus before storing at -80°C until use.

#### Pseudovirus neutralization assays

Each SARS-CoV-2 pseudovirus was titrated to standardize viral infectious dose before use in neutralization assays. Seven serial dilution of monoclonal antibodies or soluble ACE2 were added in 96-well plates, starting at 10 µg/mL for antibodies and 3 µg/mL for ACE2. As previously reported, we used soluble chimeric human ACE2 for ACE2 inhibition assays which contain ACE2 residues 1-732 fused to human IgG1 Fc. Then, pseudoviruses were added and incubated at 37 ℃ for 1 hour. In each plate, wells containing only pseudoviruses were added as controls. Vero-E6 cells were then added at a density of 4 x 10^4^ cells per well and incubated at 37 ℃ for an additional 16 hours. Cells were lysed and luminescence was determined by the Luciferase Assay System (Promega) and Tecan Infinite 200 PRO using i-control software v.3.9.1.0, in accordance with the manufacturer’s instructions. Data was analyzed in GraphPad Prism v.11 to identify the Neutralization IC50 by fitting a five-parameter dose-response curve. For fold-change calculations, IC50 values <0.002 µg/mL were assigned 0.002 µg/mL and values >10 µg/mL were assigned 10 µg/mL; fold changes involving assay-bound values are therefore reported as lower/upper bounds as appropriate.

### QUANTIFICATION AND STATISTICAL ANAYLYSIS

Neutralization IC50 values were determined by fitting a five-parameter dose-response curve in GraphPad Prism v11.

## Author Contributions

The study was conceptualized by Y.G., and D.D.H. Experiments were conducted and data analyzed by M.W., H.H., S.C., J.Y., I.A.M., C.Y.H., Q.W., Y.H., Y.G. The manuscript was written by M.W., H.H., Y.G., and D.D.H. All contributing authors have reviewed and endorsed the manuscript.

## Acknowledgements

This study was supported by funding from the NIH SARS-CoV-2 Assessment of Viral Evolution (SAVE) Program (subcontract no.0258-A700-4609 under federal contract no.75N93021C00014) and the Gates Foundation (INV019355) to D.D.H., internal startup funding UR014016 from Columbia University to Y.G., and donations from Andrew and Peggy Cherng. We thank all who contributed their data to the Global Initiative on Sharing All Influenza Data (GISAID).

---

**Fig. S1.**
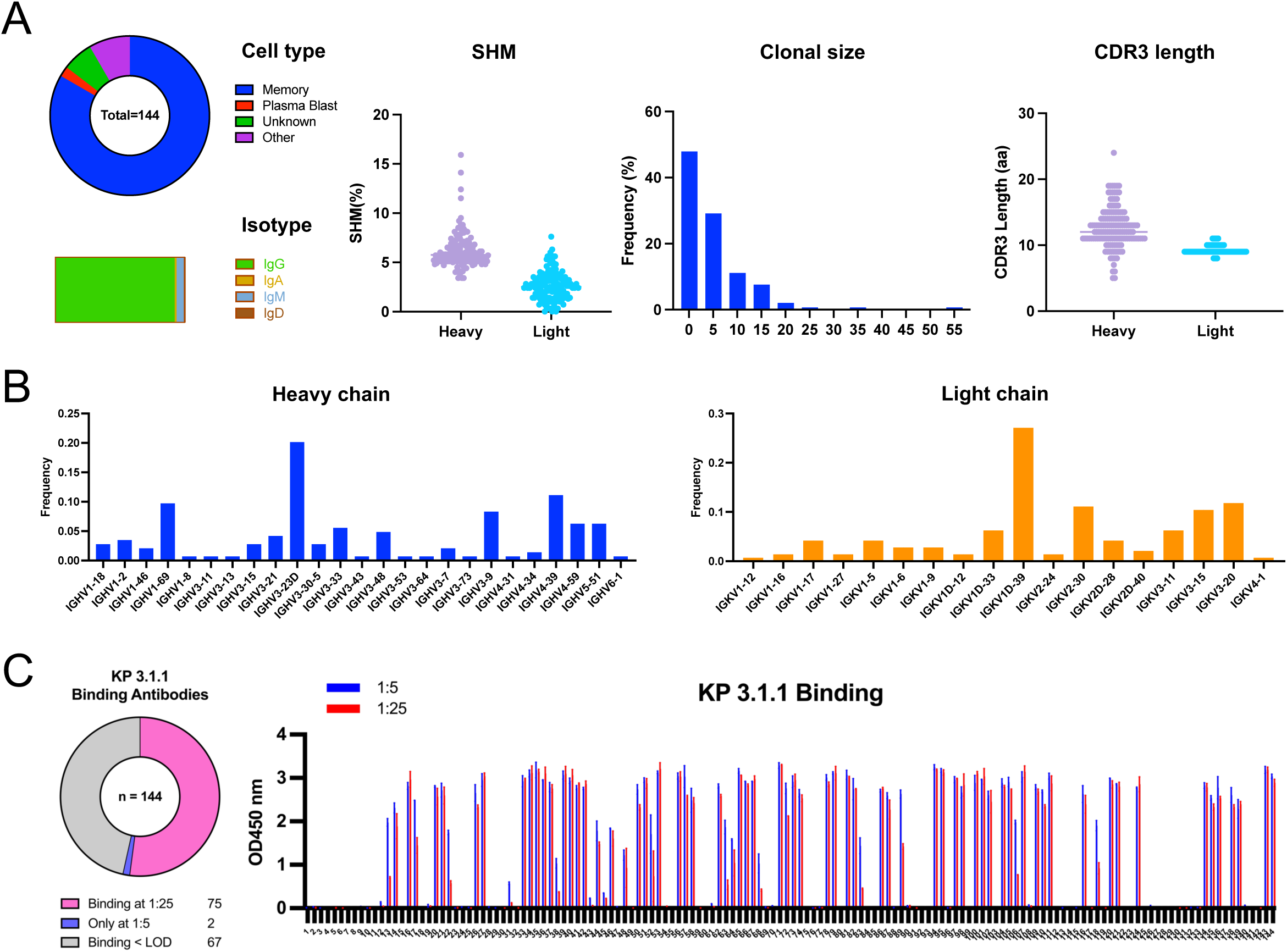
Statistics and ELISA binding of selected 144 mAbs. A. BCR features for antibodies selected from 10x genomics sequencing. B. Gene usage for selected mAbs. C. Supernatant ELISA screening of selected monocloanl antibodies against KP.3.1.1 spike.

**Fig. S2.**
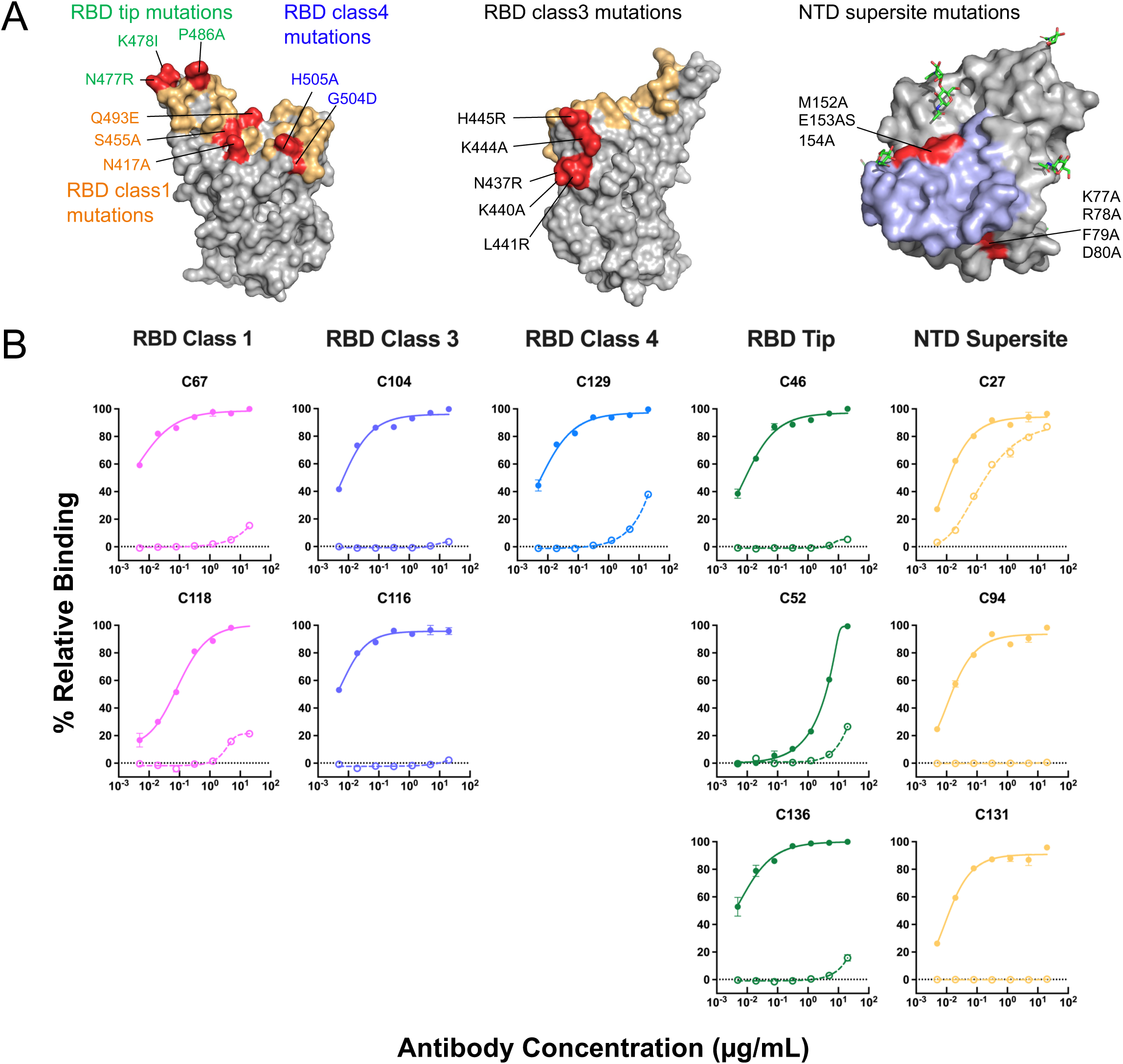
Epitope-directed mutagenesis of KP.3.1.1 spike. A. Locations of mutations used to probe the RBD class 1, RBD class 3, RBD class 4, RBD-tip, and NTD-supersite epitopes. The hACE2-binding site is shown in light orange, and the NTD supersite is shown in light blue. B. ELISA binding curves of each mAb to the wildtype KP.3.1.1 spike is denoted in filled circles, while binding curves to their corresponding epitope-mutated spike is shown in empty circles. Data are shown as mean ± standard error of mean (s.e.m.) for three technical replicates.

**Fig. S3.**
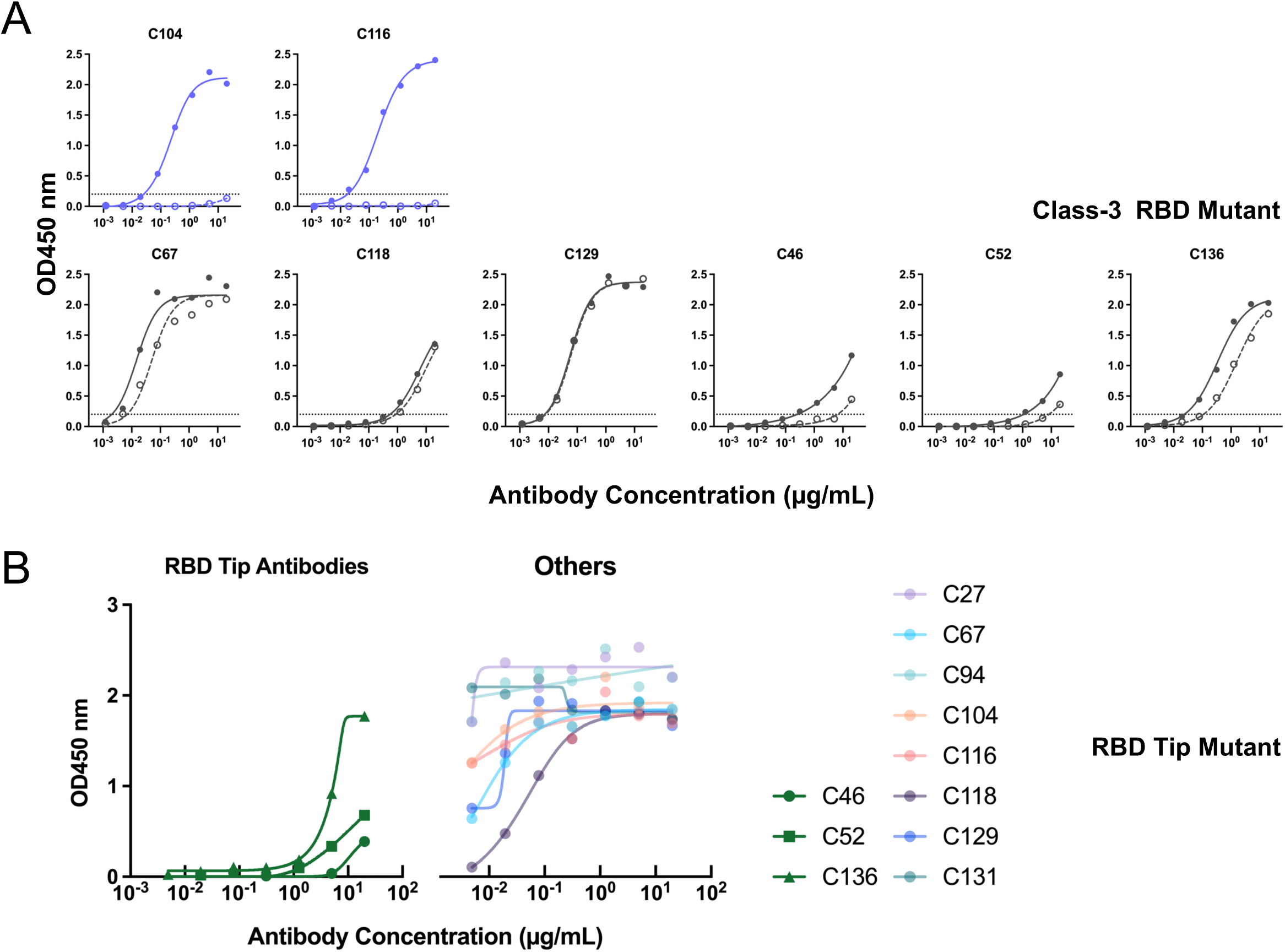
Cross-testing of epitope-directed mutants with other antibodies. A. ELISA binding curves of each mAb to the wildtype KP.3.1.1 RBD is denoted in filled circles, while binding curves to class 3 epitope-mutated RBD is shown in empty circles. B. ELISA binding of RBD tip antibodies and other C-series antibodies to RBD tip epitope-mutated spike

**Fig. S4.**
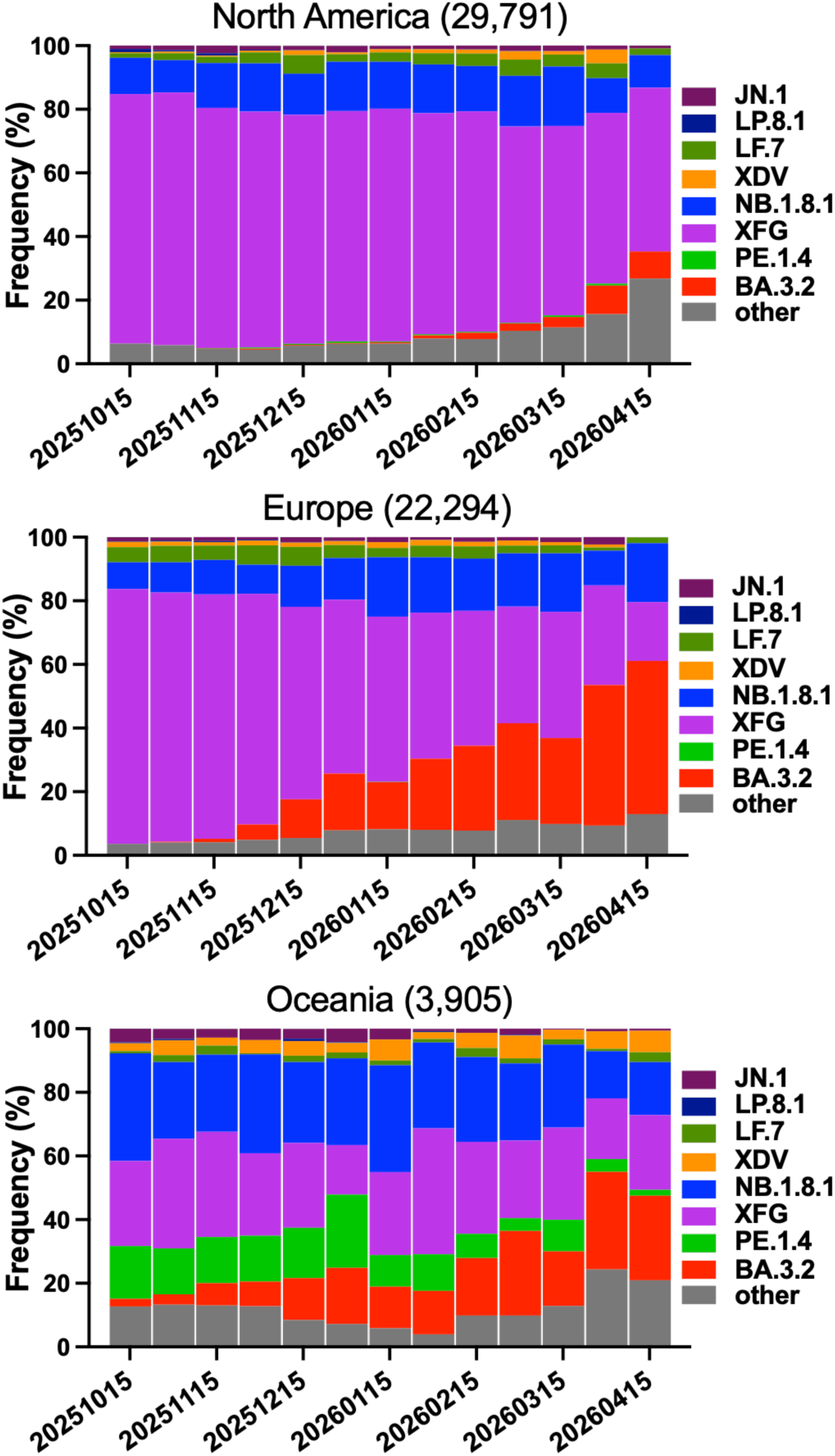
Frequencies of SARS-CoV-2 variants deposited in GISAID. The value in the upper middle of each box denotes the cumulative number of sequences for all circulating viruses in the denoted time period.

**Fig. S5.**
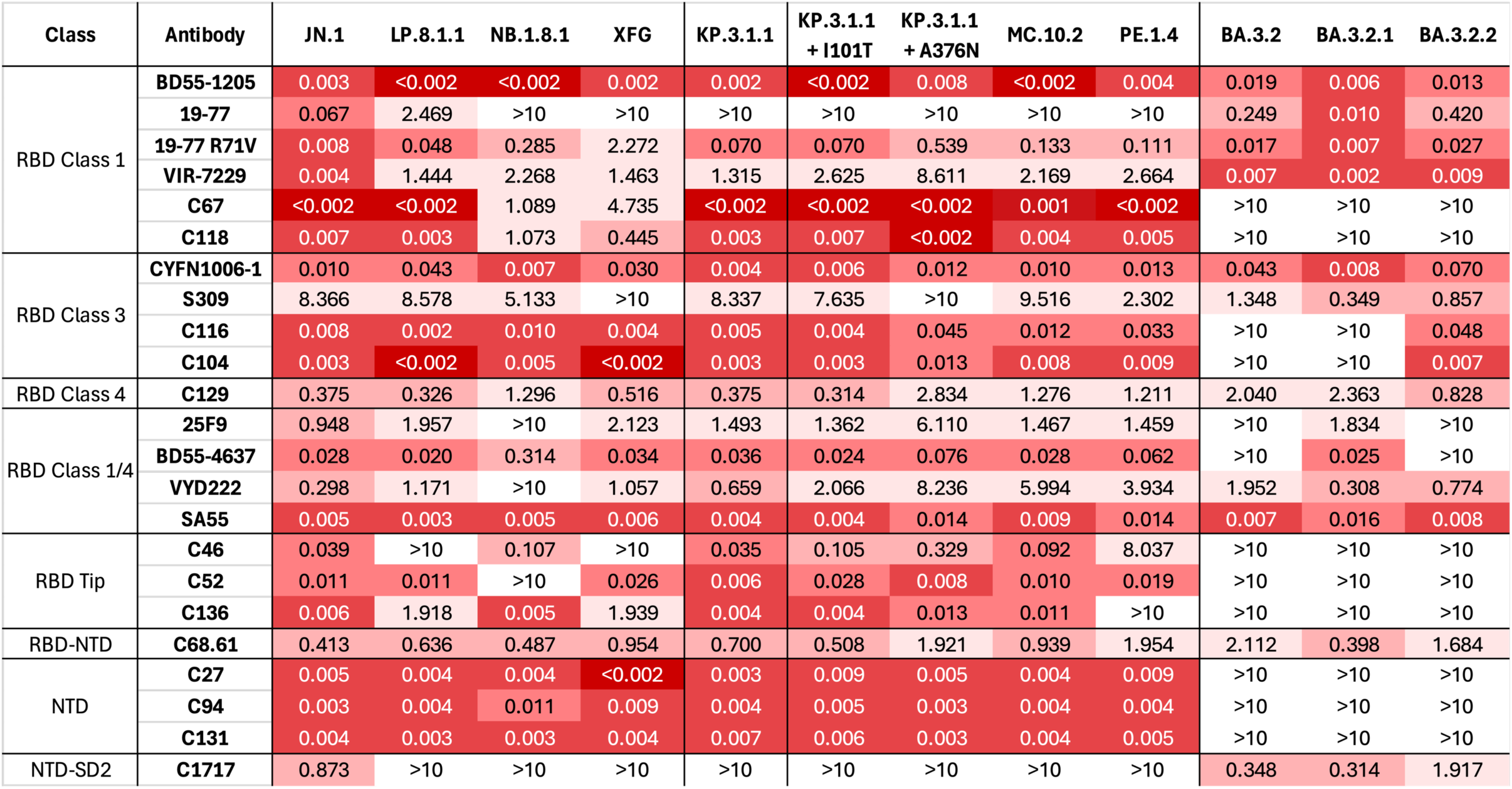
Neutralization IC50 values of the mAb panel against recently circulating SARS-CoV-2 variants.

**Fig. S6.**
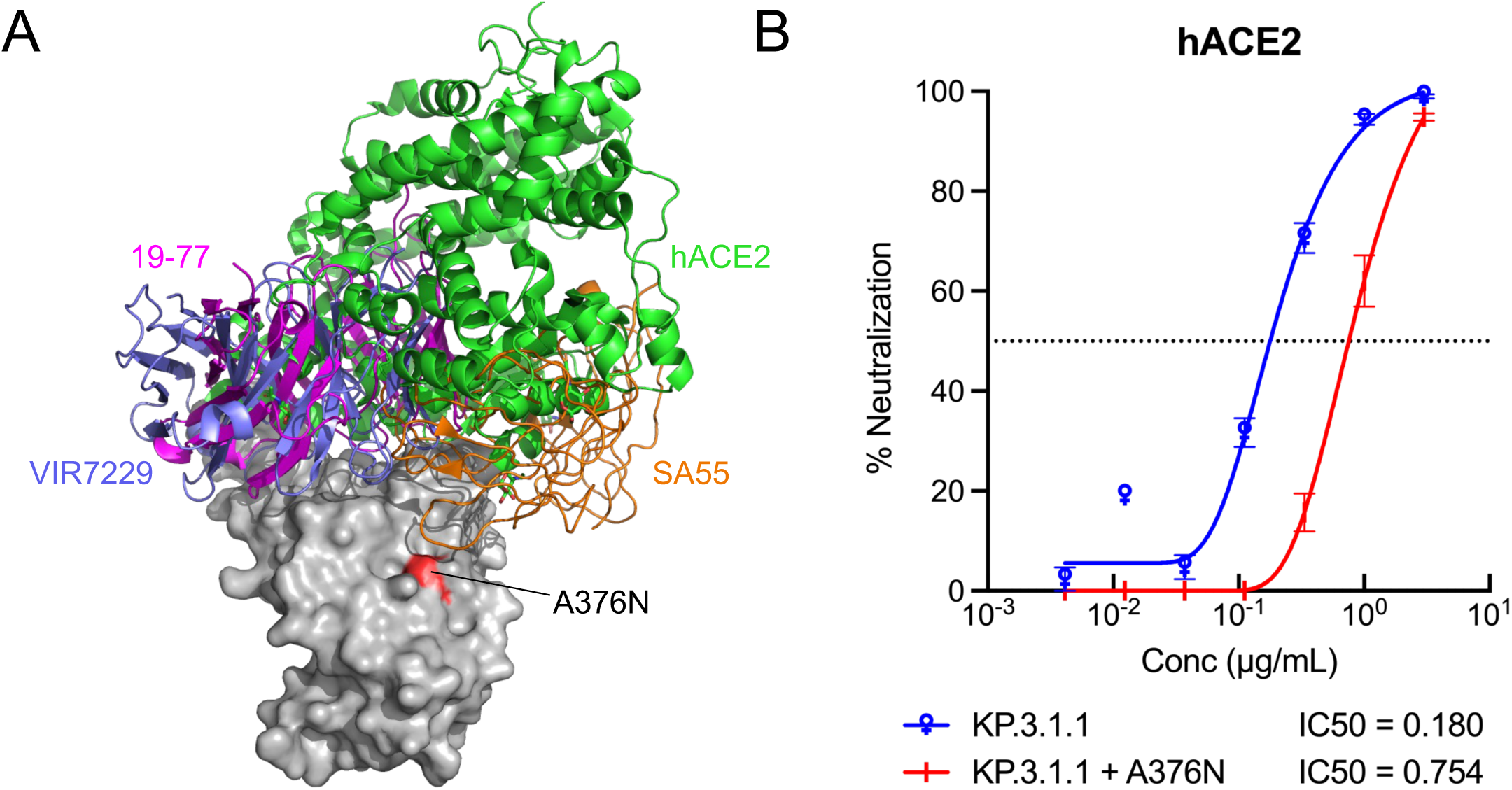
Structural context of the A376N substitution and its effect on hACE2-mediated inhibition. **(A)** Location of A376N with hACE2 and other broadly neutralizing mAbs. **(B)** Sensitivity of pseudotyped KP.3.1.1 and KP.3.1.1 with A376N to hACE2 inhibition. The hACE2 concentrations resulting in 50% inhibition of infectivity (IC50) are presented. Data are shown as mean ± standard error of mean (s.e.m.) for three technical replicates.

